# Symbiotic cyanobacteria communities in hornworts across time, space, and host species

**DOI:** 10.1101/2020.06.18.160382

**Authors:** Jessica M. Nelson, Duncan A. Hauser, Fay-Wei Li

## Abstract

**Rationale:** While plant-microbe interactions have been intensively studied in mycorrhizal and rhizobial symbioses, much less is known about plant symbioses with nitrogen-fixing cyanobacteria. Here we focused on hornworts (a bryophyte lineage), and investigated the diversity of their cyanobionts and how these communities are shaped by spatial, temporal, and host factors.

**Method:** We carried out repeated samplings of hornwort and soil samples in upstate New York throughout the growing season. Three sympatric hornwort species were included, allowing us to directly compare partner specificity and selectivity. To profile cyanobacteria communities, we established a new metabarcoding protocol targeting *rbcL-X* with PacBio long reads.

**Results:** Hornwort cyanobionts have a high phylogenetic diversity, including clades that do not contain other known plant or lichen symbionts. While the sympatric hornwort species have similarly low specificity, they exhibit different preferences toward cyanobionts, although this depended on what cyanobacteria were present in the soil. Cyanobacterial communities varied spatially, even at small scales, but time did not play a major organizing role.

**Conclusion.:** This study highlights the importance of sampling soil and sympatric species to infer partner compatibility and preference, and marks a critical step toward better understanding the ecology and evolution of plant-cyanobacteria symbiosis.

## Introduction

Symbiotic interactions with microbes are key drivers in plant evolution and ecology. They not only facilitated plants’ initial transition to land (Bidartondo *et al*., 2011), but also played a pivotal role in subsequent species diversification and ecosystem functioning (Lugtenberg & Kamilova, 2009; Van Der Heijden *et al*., 2015; Lutzoni *et al*., 2018). The most well-studied plant-microbe mutualisms are those with arbuscular mycorrhizal fungi (AMF) and rhizobia. In both cases the symbiosis is endophytic and plants provide carbon sources to the symbionts in exchange for phosphate (AMF) or nitrogen (rhizobia and AMF). The natural histories and genetic mechanisms for these two types of symbiosis have been intensively studied (Oldroyd *et al*., 2011; Van Der Heijden *et al*., 2015; Andrews & Andrews, 2017; MacLean *et al*., 2017).

Cyanobacteria also form specialized nutritional symbioses with plants in which they exchange fixed nitrogen for carbon supplements (Rai *et al*., 2000), but these relationships have been much less studied than those with AMF and rhizobia. Five disparate plant lineages have independently evolved the ability to engage in this specialized symbiosis with cyanobacteria: the angiosperm genus *Gunnera*, all cycads (gymnosperm), the water fern genus *Azolla*, a small family of liverworts (Blasiaceae), and all hornworts (Meeks, 1998; Rai *et al*., 2000). In all these lineages, the symbiotic cyanobacteria (hereafter cyanobionts) are hosted inside specialized plant structures that develop independent of contact with cyanobacteria, for example slime cavities in hornworts and coralloid roots in cycads (Rai *et al*., 2000; Santi *et al*., 2013). The cyanobionts are all extracellular endophytes except in *Gunnera* where they are intracellular in the cells of the stem glands (Rai *et al*., 2000). Research on plant-cyanobacteria symbiosis mechanisms thus far has determined the sequence of events for establishment of the interaction and some of the necessary cyanobacterial genes (Santi *et al*., 2013). The symbioses have been experimentally reconstituted in hornworts, liverworts, *Gunnera*, and cycads (Enderlin & Meeks, 1983; Ow *et al*., 1999; Chiu *et al*., 2005; Santi *et al*., 2013).

Plant cyanobionts belong to cyanobacterial groups able to produce nitrogen-fixing heterocysts and motile hormogonia (Santi *et al*., 2013). Most of these plant cyanobionts have been commonly referred to as *Nostoc* (Rai et al., 2000), but this genus is notoriously polyphyletic and contains multiple distinct genetic clades (Rajaniemi *et al*., 2005; Shih *et al*., 2013; Komárek, 2016). The taxonomic confusion for *Nostoc* could conceal wider symbiont diversity and patterns of compatibility and preference between partners. Until recently, research on the diversity of plant cyanobionts had relied on culturing the cyanobacteria or limited methods of genetic differentiation such as RFLPs (Leizerovich *et al*., 1990; West & Adams, 1997; Costa *et al*., 1999; Costa *et al*., 2001; Guevara *et al*., 2002; Zheng *et al*., 2002; Papaefthimiou *et al*., 2008; Rikkinen & Virtanen, 2008; Thajuddin *et al*., 2010; Yamada *et al*., 2012; Liaimer *et al*., 2016). Metabarcoding and metagenomic methods using next generation sequencing platforms have begun to be applied recently, covering *Azolla*, cycads, and one hornwort (Dijkhuizen *et al*., 2018; Zheng *et al*., 2018; Suárez-Moo *et al*., 2019; Zheng & Gong, 2019; Bell-Doyon *et al*., 2020; Bouchard *et al*., 2020).

These next generation sequencing studies have relied on rRNA regions for identification of bacteria, but this approach poses several challenges in characterizing symbiotic cyanobacteria communities. First, the standard 16S primers are nonspecific and will amplify the entire bacterial community as well as plant organelles, limiting the measurement of cyanobacteria in a given sample. This is a particular challenge when targeting cyanobacteria since they share ancestry with plastids. Second, the short 16S amplicons obtained with Illumina sequencing do not provide much phylogenetic signal. This is especially problematic because the classification of cyanobacteria—from order to species levels—has been largely unresolved, so assigning accurate taxonomy with conventional 16S classifiers is difficult. Third, cyanobacterial genomes typically contain multiple rRNA operons which skews read counts, and in some cases the operons can be highly divergent in the same genome (Johansen *et al*., 2017), which makes rRNA genes inappropriate species markers in this case.

Concomitant with these challenges of defining and surveying cyanobacterial diversity is a lack of information on the environmental factors controlling the composition of cyanobiont communities. Only a few studies have attempted to survey the cyanobacterial diversity of soils near hosts (West & Adams, 1997; Cuddy *et al*., 2012; Liaimer *et al*., 2016; Suárez-Moo *et al*., 2019) and have found differing amounts of overlap between soil and symbiotic communities. It is also unclear how geographic distance and host species affect the communities. While some studies have reported very little overlap of cyanobacterial taxa between sites (West & Adams, 1997; Bouchard *et al*., 2020), others have reported the appearance of the same symbiont on distant continents (Rikkinen & Virtanen, 2008; Gehringer *et al*., 2010). Studies that have compared different host species so far have not detected clear specificity of symbionts to certain hosts (Gehringer *et al*., 2010; Suárez-Moo *et al*., 2019), but this may be due in part to host species and location being confounded. In addition, none of the previous studies have explicitly tested the effects of time, a commonly ignored dimension in plant microbiome studies in general (Peršoh, 2015; Wagner *et al*., 2016). Plants’ symbiotic structures are usually continuously developed and colonized by cyanobacteria (Adams & Duggan, 2008), which implies that cyanobiont compositions may be dynamic over time.

Out of the five plant groups that host cyanobionts, hornworts are particularly well suited to investigate the ecological and evolutionary dynamics of cyanobiont communities. They are more broadly distributed globally than the other cyanobacterial host groups. They are also more speciose and have deeper evolutionary splits (Villarreal *et al*., 2015) than all the host groups besides cycads. These features present the opportunity for biogeographic studies across biomes as well as macroevolutionary studies across geological time. Furthermore, hornworts are much more tractable laboratory models than the other groups. Hornworts can be grown in axenic culture for reconstitution experiments more quickly and easily and experiments can use identical clones rather than needing to germinate many genetically distinct seeds to serve as experimental replicates. Indeed, the hornwort *Anthoceros punctatus* and cyanobacterial strain *Nostoc punctiforme* PCC 73102 have long been used to investigate symbiotic physiology (Meeks, 2003). Finally, several high-quality hornwort and cyanobiont genomes have been recently published, accelerating research on the genetic mechanisms of the symbiosis (Nelson *et al*., 2019; Zhang *et al*., 2020; Li *et al*., 2020).

In this study, we address a number of the existing knowledge gaps in the diversity and ecology of cyanobacterial symbiosis with a survey of hornwort cyanobionts. We profile the cyanobacterial communities with PacBio circular-consensus sequencing (CCS) of full length *rbcL-X* amplicons. Our sampling of three sympatric hornwort species and adjacent soils across the hornwort growing season allows us to investigate not only the phylogenetic diversity of hornwort cyanobionts, but also the effects of host species, various spatial scales, and time on cyanobiont community compositions.

## Materials and Methods

### Sample collection

Hornworts and adjacent soil were sampled at two sites in Tompkins County, New York about 20 km apart: one in the Cornell Botanic Gardens on the bank of Grossman Pond and one on a roadside bank in Potato Hill State Forest (Fig. 1**a, b**). The Grossman Pond site had a population of *Notothylas orbicularis* while the Potato Hill site had a mixed population of *N. orbicularis, Phaeoceros carolinanus*, and *Anthoceros agrestis* (Fig. 1**c**). These three species are the most common, if not the only, hornworts in the Northeastern USA (Schuster, 1992).

**Figure 1.**
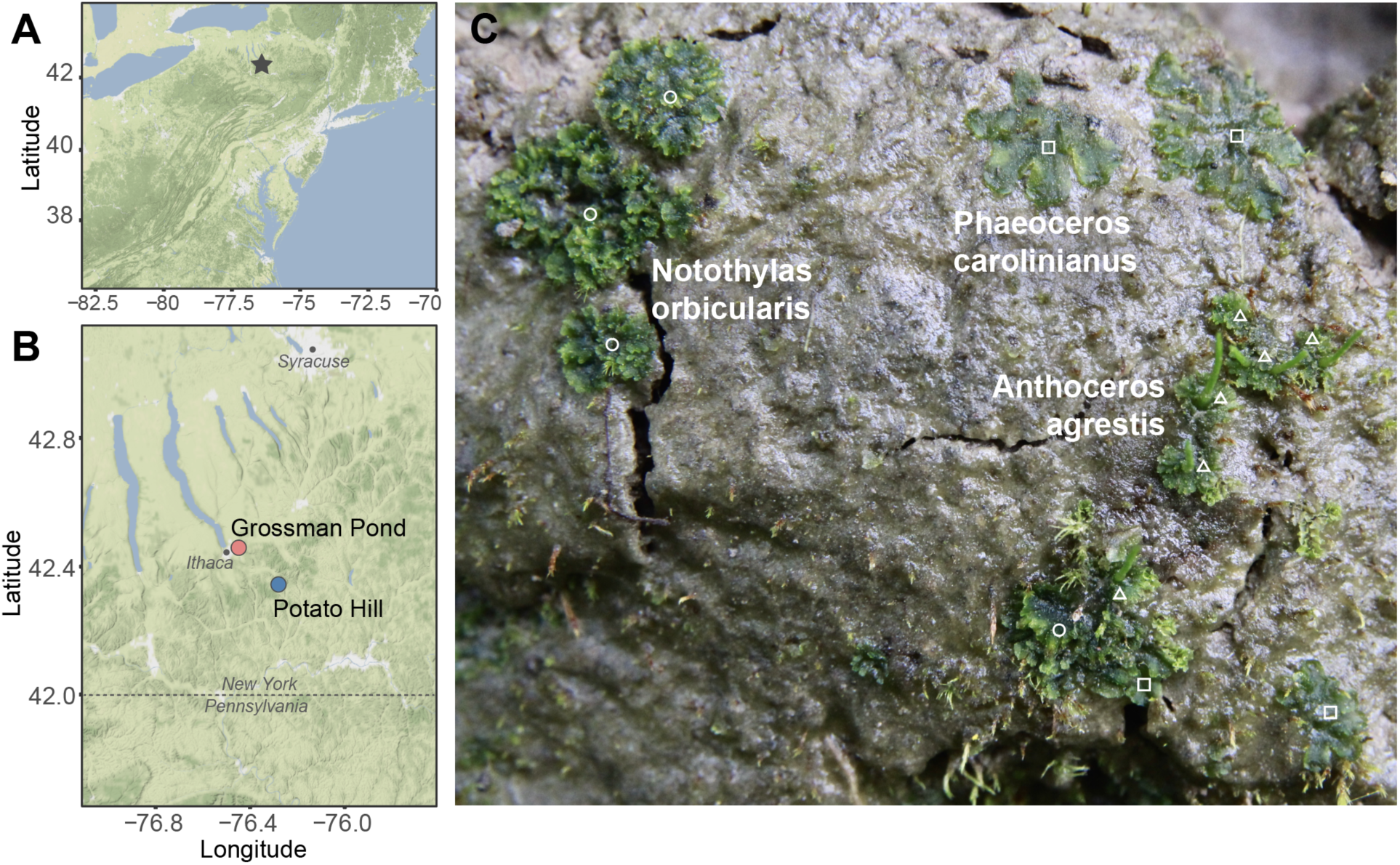
Hornwort population samplings. (A) Grossman Pond and Potato Hill sites are located in upstate New York (star). (B) A map showing the locations of the two sites. (C) Three sympatric hornwort species can be found growing together at the Potato Hill site. Circle: *Notothylas orbicularis*. Square: *Phaeoceros carolinianus*. Triangle: *Anthoceros agrestis*.

In the study area, hornworts are annual with a growing season typically starting in early- to mid-summer and ending in fall. To capture variation over this growing season, samples were taken every three weeks from August 23 to October 30 in 2018, for a total of 4 sampling times at each site.

Quadrats of 0.25 m^2^ were used as the sampling unit to delineate patches of hornworts to sample and pin flags were used to mark the locations so the same ground area was used each time. Two quadrat locations (about 20m apart) were sampled at Grossman Pond and three at Potato Hill (about 20 and 7 m apart). Within each quadrat, four soil samples were collected along the diagonals using sterile 15 mL falcon tubes. Four plugs of soil approximately one cm deep were taken for each sample, moving from the center of the quadrat towards one corner. Four plants (gametophytes, sometimes with attached sporophytes) of each present hornwort species were collected within the area of the quadrat using an ethanol-cleaned knife. *Notothylas* declined toward the end of the season earlier than the other species, resulting in two quadrats at the final time point lacking this species. Soil and plant samples were stored at 4? until processing (usually about one day, but no longer than 48 hours).

### Soil sample preparation

A total of 76 soil samples were collected. Soils were passed through a clean 2mm mesh sieve to even out particle sizes, remove large debris or organic materials, and mix the sample together. From this mixture, a small lump of soil (about 150-250 mg) was put directly into an E.Z.N.A. soil DNA kit (Omega Bio-Tek) disruptor tube with glass beads. Between samples, adhering soil was washed off of the sieve and the sieve was cleaned with 70% ethanol. DNA was extracted from soils using the E.Z.N.A. soil DNA kit (Omega Bio-Tek) with two cHTR reagent washes.

### Hornwort sample preparation

A total of 168 hornwort samples were collected. Fresh hornworts were cleaned in sterile water to remove all adhering soil. They were first rinsed in multiple changes of sterile water using vortexing and forceps to dislodge soil particles. Once most of the soil was removed, the plants were sonicated in sterile water at 1% amplitude for 15 seconds in intervals of 3 seconds with 2 second rests in between. This protocol was determined by testing for the strongest sonication that would not break the hornwort cells. Plants were then blotted dry on clean Kimwipes and transferred to homogenization tubes with 2mm zirconia beads. Tubes holding the plants were frozen with liquid nitrogen and ground for two minutes at 1,300 strokes/min on a SPEX SamplePrep 1600 MiniG tissue homogenizer in a metal box pre-chilled in liquid nitrogen. DNA was extracted from plant tissues using the E.Z.N.A. Plant DNA Kit (Omega Bio-Tek).

### Amplicon library preparation and sequencing

We targeted the cyanobacteria-specific single copy *rbcL-X* region (∼800bp) and used the PacBio CCS method to sequence the long amplicons. The *rbcL-X* region was amplified by PCR from all DNA samples using barcoded versions of the primers CW and CX (Rudi *et al*., 1998). Eight forward primer barcodes and twelve reverse primer barcodes were used to make 96 dual barcode combinations (Table S1). Samples were amplified with the appropriate barcoded primers using Phusion high-fidelity polymerase (New England Biolabs). The PCR recipe and thermocycler program can be found in Table S2.

A mock community of five cyanobacteria was used as a positive sequencing control. DNA was extracted from pure laboratory cultures of cyanobacteria with a standard CTAB protocol and grinding with copper beads, as previously described (Nelson *et al*., 2019). Then *rbcL-X* was amplified from each using the same PCR program as above with unbarcoded primers. The concentrations of these PCR products were measured with the Qubit HS DNA kit and mixed in equal quantity. This mix was then used in barcoded reactions in the same way as all other samples.

The amplicon concentrations were measured by the Qubit HS DNA kit, and samples pooled in equal quantity into 3 libraries. Libraries were cleaned using ProNex beads (Promega) with a 1.5:1 ratio of beads to sample. PacBio circular consensus sequencing was done on the Sequel platform with v3.0 chemistry at Duke University. The sequencing reads were deposited at NCBI SRA under the accession PRJNA632853.

### Data processing

We used the PacBio ccs (v4.0.0; https://github.com/PacificBiosciences/ccs) package to generate circular consensus sequences from the raw PacBio reads, and only kept those with at least five complete passes and a predicted consensus accuracy over 0.999 (--min-passes=5 --min-rq=0.999). Demultiplexing was done by lima (v1.10.0; https://github.com/PacificBiosciences/barcoding) with the minimum barcode score set to 26 (-- min-score 26), followed by primer removal and read reorientation by the removePrimers function of the dada2 package (Callahan *et al*., 2016). We further filtered the dataset using the dada2 filterAndTrim function to remove sequences longer than 1200bp, shorter than 400bp, or having a quality score lower than 3 (minQ=3, minLen=400, maxLen=1200). Dereplication, error learning, amplicon sequence variant (ASV) inference, and chimera detection were done with the newly developed dada2 PacBio pipeline (Callahan *et al*., 2019). In contrast to the traditional clustering-based method, which uses an arbitrary identity cutoff to group sequences into operational taxonomic units (OTUs), the ASV approach incorporates models of sequencing error to derive exact sequence variants. To benchmark the ASV approach against traditional OTU clustering, we used VSEARCH (Rognes *et al*., 2016) to carry out dereplication, chimera removal, and clustering at 97% and 95% sequence identities. Resulting ASV and OTU tables were analyzed using the phyloseq (v1.30.0) R package (McMurdie & Holmes, 2013).

### Phylogenetic analysis

Phylogenies were inferred using two datasets. The first dataset includes ASVs from this study and a large collection of cyanobacterial *rbcL-X* sequences, so that we can place our ASVs onto a broader phylogenetic context. To build a *rbcL-X* database, we (1) downloaded all the available cyanobacterial genomes and extracted their *rbcL-X* region, (2) queried all the ASVs by BLASTn against the NCBI database and retrieved the hits (with an e-value threshold of 0.001), and (3) incorporated *rbcL-X* sequences (done by Sanger sequencing) from some of our own cyanobacterial isolates. Redundant and identical sequences were removed. Combining ASVs from the present study and previously sequenced references resulted in a total of 1748 sequences. These were aligned using PASTA (Mirarab *et al*., 2015). The phylogeny was inferred using RAxML-HPC2 v8.2.12 (Stamatakis, 2014) on CIPRES (Miller *et al*., 2010), with 1000 replicates of rapid bootstrapping to assess branch supports.

In order to better visualize and delineate the hornwort cyanobiont clades, we analyzed a second dataset including only the ASVs. For the plant samples in this dataset, we filtered out low abundance ASVs (<3% per sample) because they likely represent epiphytic or soil bacteria, or PCR/sequencing errors. This threshold was selected based on the mock community results (see below). The alignment was done by PASTA and tree inference by RAxML as before. We plotted the number of times an ASV can be found in each sample category (soil, *Anthoceros, Notothylas*, and *Phaeoceros*) using ggtree (Yu *et al*., 2018). We defined cyanobiont clades as those containing ASVs present in at least 3 plant samples and with bootstrap support over 95. To assign taxonomic affinities to nodes, we cross-referenced this phylogeny with (1) the one described above with other known cyanobacterial sequences, and (2) trees from Shih et al. (2013) based on a large sampling of cyanobacteria genomes, and from Otalora et al. (2010) and Magain et al. (2017; 2018) on lichen photobionts. Due to the relatively poor taxonomic understanding of cyanobacteria, we cannot confidently assign families to nodes. Instead we listed, if present, the strains/isolates that are present in the same clade as each ASV group.

### Analysis of cyanobacteria diversity and community composition

We used the R package phyloseq (v1.30.0) (McMurdie & Holmes, 2013) to transform ASV counts per sample into relative abundance, calculate Unifrac and weighted Unifrac distance between samples, and carry out a series of principal coordinate analyses (PCoA). To determine if cyanobacteria communities significantly vary by sampling location, time, and host species, PERMANOVA tests were done with 10,000 permutations using the adonis function in the vegan package (Dixon, 2003). To identify specific ASVs that show differential abundance among host taxa, Analysis of Composition of Microbiomes (ANCOM) (Mandal *et al*., 2015) was conducted in the R package ANCOM (v2.1; https://github.com/FrederickHuangLin/ANCOM). We carried out the PERMANOVA and ANCOM analyses with low abundance ASVs (3%) removed.

Chao, Shannon, and Simpson alpha diversity indices were calculated by phyloseq, and Faith’s Phylogenetic Diversity and Mean Pairwise Distance indices was calculated by the R package picante (Kembel *et al*., 2010) with the ASV-only phylogeny from above. The UpSetR package was used to visualize the overlaps of ASVs among quadrats, sample types, and time points (Conway *et al*., 2017). The scripts, commands, and data matrices from this study can be found at https://github.com/fayweili/hornwort_cyano_interaction.

Based on the community compositions, we assessed specificity and selectivity of hornwort-cyanobiont combinations. We follow the definitions of Bubrick et al. (1985) and define specificity as partner compatibility and selectivity as partner preference.

## Results

### Amplicon sequencing

A total of 288 samples were sequenced on three PacBio Sequel runs, 244 of which were from the focal Grossman Pond and Potato Hill samplings (the rest were controls and other samples not part of this study). Over the three PacBio runs, we obtained 1,573,867 raw sequences, which yielded 669,780 CCS reads passing filters on number of passes, accuracy, length, and correct barcodes and primers; of these, 598,442 reads were from the focal samplings.

Our benchmark on mock communities clearly demonstrated that the ASV approach has a much higher accuracy and sensitivity than traditional OTU clustering (Fig. 2). The mock community consisted of five cyanobacteria strains and was included in each PacBio sequencing run. In two out of the three mock samples, exactly five ASVs were recovered, and their sequences were identical to those from the input strains (Fig. 2**d**). One mock sample returned one extra ASV (ASV25), which was not part of the mock community. However, this ASV has a low relative abundance (3%; Fig. 2**e**). In mock 1 and 3, the ASV counts significantly deviated from equal abundance (Chi-square *p*=0.002 and 0.047, respectively; Fig. 2**e**), but this could be due to an extra PCR step that was used when making the mock, potentially introducing more biases. No significant count difference was detected in mock 2 (*p*=0.815; Fig. 2**e**). In contrast, the OTU method recovered many more taxa than were actually included (12 and 7 OTUs on average per mock when clustered at 97% and 95%, respectively, after removing singletons; Fig. 2**f**,**g**).

**Figure 2.**
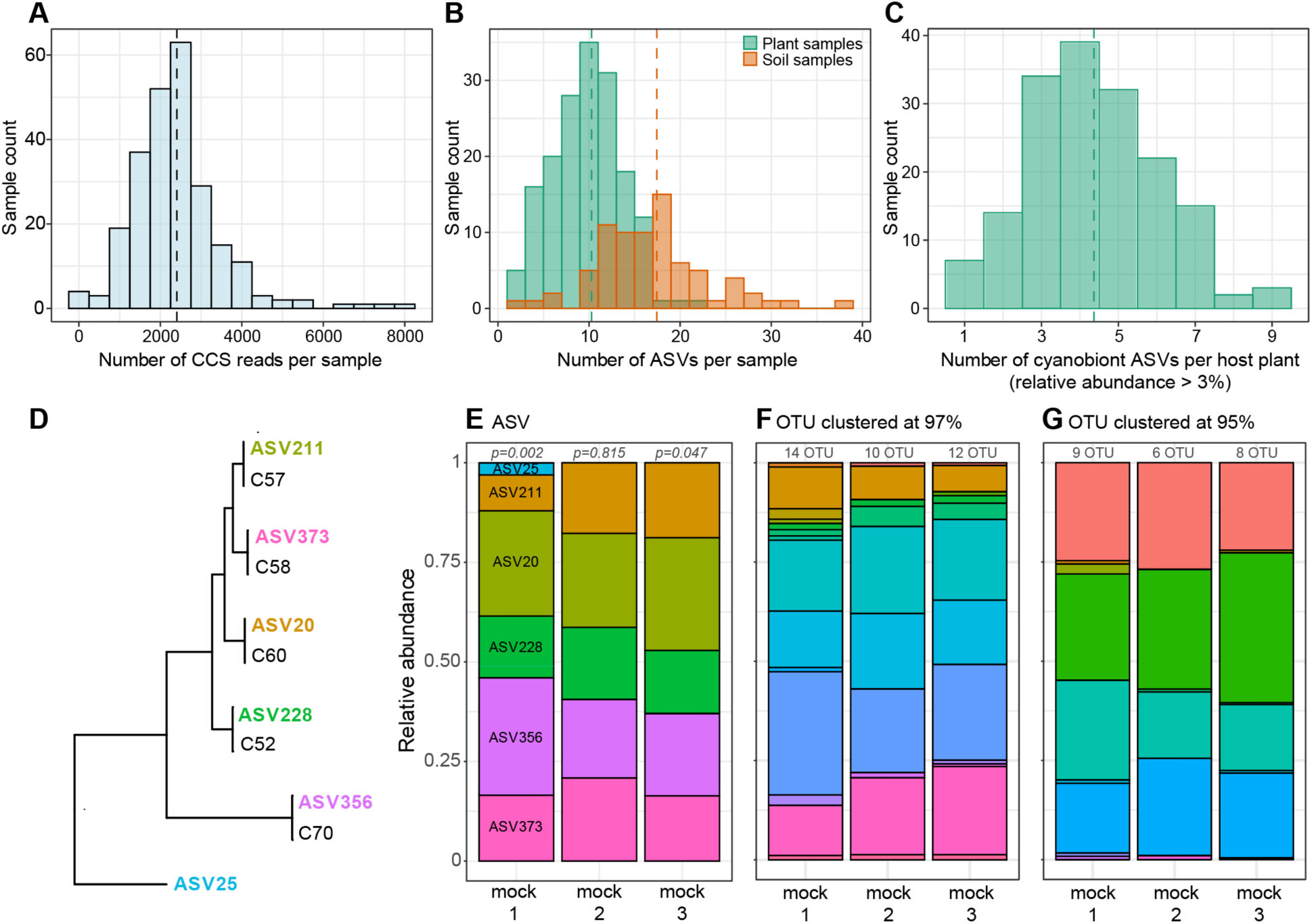
Summary of sequencing results. Distributions of (A) final filtered CCS reads, (B) all ASVs, and (C) ASVs from plants with relative abundance over 3%. Dashed lines are the group means. (D) The mock community consists of five isolates, and was sequenced in three separate PacBio runs (mock 1-3). A total of 6 ASVs were recovered from mock1-3, five of which have identical sequences with the original isolates. The only exception is ASV25, which was not part of the mock community and only found in mock 1 and in low abundance. (E) Relative abundance of ASVs in mock1-3. The italic numbers above bars are chi-square *p* values tested against equal abundance. OTU clustering-based approach at (F) 97% and (G) 95% performed poorly in our PacBio datasets and cannot accurately recover the mock strains.

In total, 382 ASVs were inferred from the soil and plant samples (Fig. 2**a-c**). The rarefaction curves from individual samples all plateaued (Fig. S1), and across all samples, the correlation between read count and ASV number is very weak (R^2^=0.0145, *p*=0.0317; Fig. S1), which suggest sufficient sequencing coverage. As expected with our sampling design of mixing multiple soil plugs together, soil samples usually harbored significantly more ASVs than plants (average ASVs/sample: 17.4 vs 10.3, *p*<10^−14^ unpaired t-test; Fig. 2**b**). After filtering out ASVs with lower than 3% relative abundance, a hornwort sample on average has about 4.4 cyanobiont ASVs (Fig. 2**c**). Over 67% of the hornwort samples contain an ASV accounting for at least 50% of the reads (Fig. S2).

### Phylogenetic diversity of hornwort cyanobionts

Because of the unresolved nature of cyanobacteria taxonomy, we used the “subclade” scheme from Shih et al. (2013) to provide a higher level classification for our ASVs. The soil cyanobacteria from our study spanned a wide phylogenetic range, including members from Subclade A (e.g. *Oscillatoria*), Subclade F (e.g. *Pseudanabaena*), and a few that cannot be mapped onto the Shih et al. (2013) phylogeny (Fig. 3, Fig. S3). All the hornwort cyanobiont ASVs fell under Subclade B1, which roughly corresponds to Subsection IV (Nostocales) plus Subsection V (Stigonematales) from Castenholz et al (2001). Subsections IV and V were circumscribed solely based on morphology, and were not recovered as monophyletic groups in our *rbcL-X* phylogeny.

**Figure 3.**
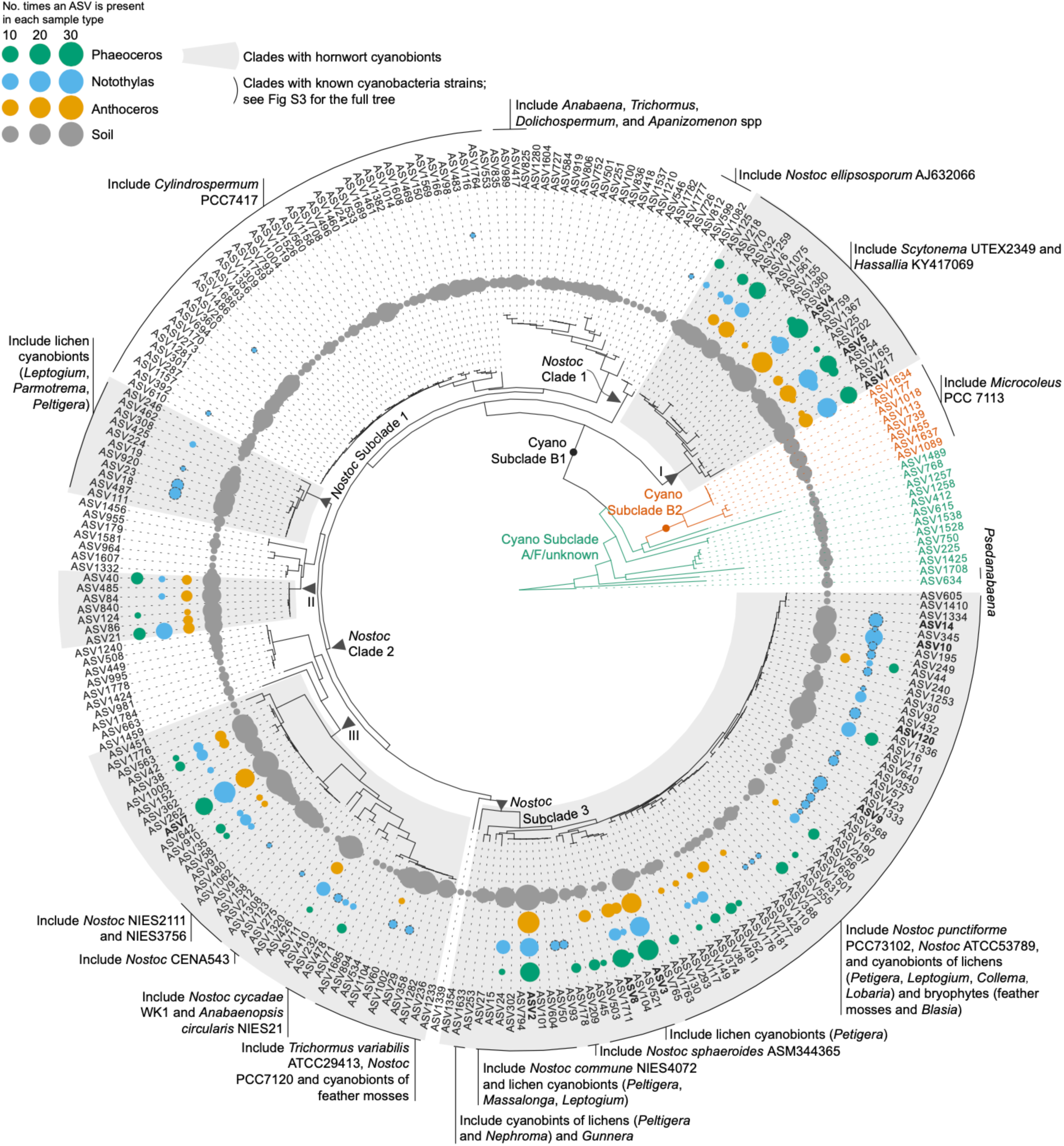
Phylogeny of cyanobacteria ASVs. The designations of “Cyano Subclades” were based on Shih et al. (2013), and “*Nostoc* Subclades” based on Otalora et al. (2010) and Magain et al. (2017, 2018). Blue circles with dashed outlines are *Notothylas* cyanobionts exclusively from Grossman Pond. ASVs mentioned in the text were in bold.

The majority of the available sequences in Subclade B1 were annotated as “*Nostoc*”, although this genus is clearly a polyphyletic assemblage. We mapped the *Nostoc* clade and subclade designations made by previous cyanolichen studies (Otálora *et al*., 2010; Magain *et al*., 2017; Magain *et al*., 2018) to our phylogeny (arrowheads in Fig. 3). While these clade labels do not cover all the lineages in B1 (Fig. 3), they nonetheless provide useful anchors for comparing across studies. Hornwort cyanobionts fall into five monophyletic groups, two of which overlap with lichen photobionts (“*Nostoc* subclade 1” and “*Nostoc* subclade 3”), and one of which includes cycad cyanobionts and cyanobacteria epiphytically associated with feather mosses (group III). The remaining two clades have no other known symbiotic members (groups I and II).

### Soil cyanobacterial pools and cyanobiont communities differ over short distances

The PCoA of all samples based on weighted unifrac distance shows two clusters of soil samples, one exclusively Potato Hill samples and the other mostly Grossman Pond ones (Fig. 4**a**), indicating phylogenetically distinct communities in the soil of the two sites. Sites formed significantly different clusters in the ordination overall (PERMANOVA R^2^=0.218, *p*<10^−4^; Table S3). While the plant samples still separate somewhat by site, they show much more variation and cluster most densely in a different corner of the ordination space from either of the soil clusters (Fig. 4**b**). Even so, there is a substantial overlap between the soils and the cyanobiont communities, with 73.9% of the hornwort ASVs also found in the soil samples (Fig. 4**d**).

**Figure 4.**
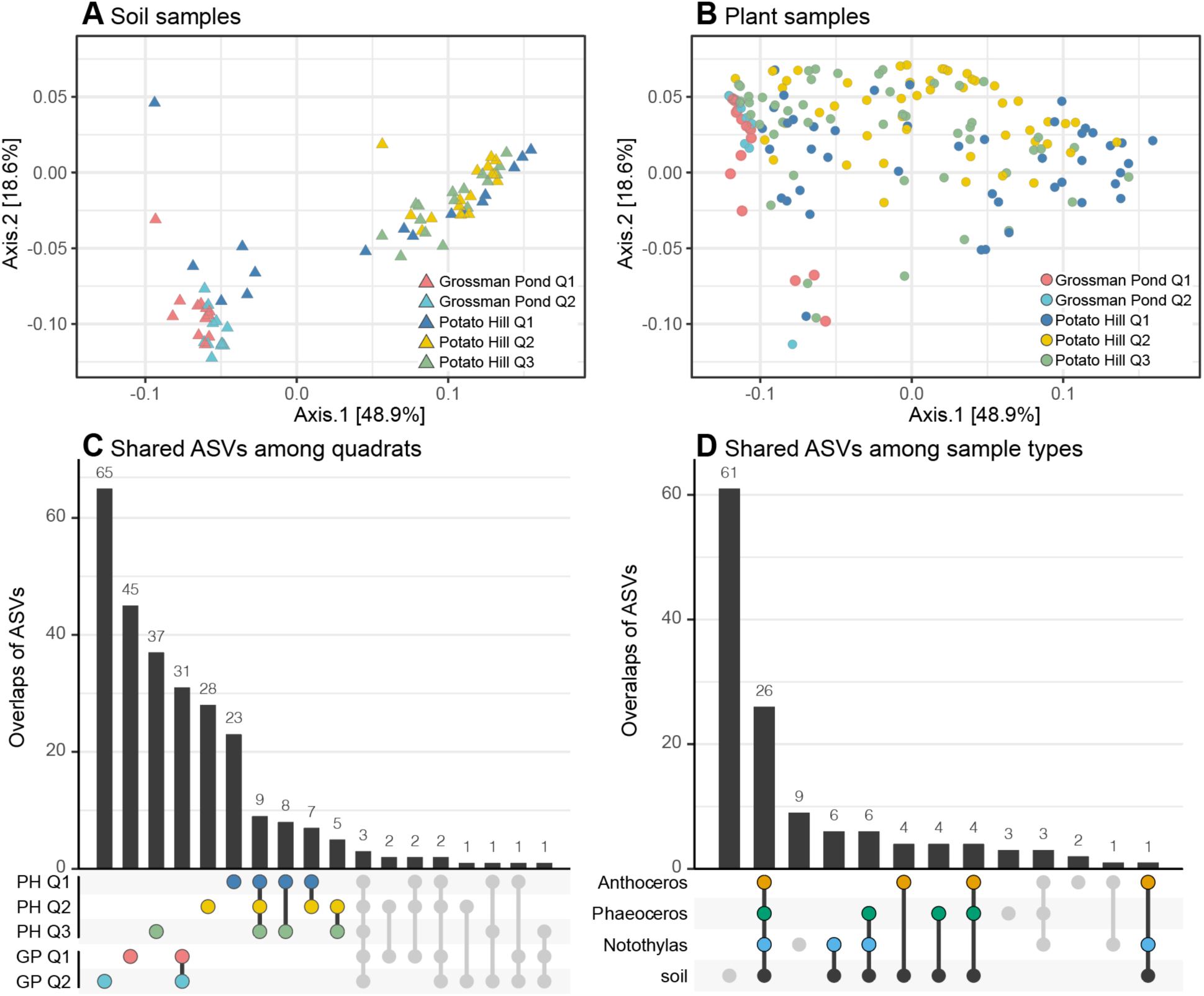
Comparison of soil and cyanobiont community composition. PCoA based on weighted unifrac distance of all samples with (A) soil samples and (B) plant samples shown on separate plots for clarity. (C) Shared and unique ASVs among the five quadrats. All the top categories are either unique or shared ASVs within the same sampling sites (highlighted in color). (D) Shared and unique ASVs among the sample types at Potato Hill site. Most cyanobiont ASVs can also be found in soil samples (highlighted in color). For (C) and (D) overlaps with zero count were not plotted. GP: Grossman Pond; PH: Potato Hill.

Samples from the same quadrat cluster together on the PCoA of Potato Hill plants with unweighted unifrac, but this structure disappears when weighted unifrac is used (Fig. S4). Similarly, soils mostly separate by quadrat using either unifrac (Fig. S4). In line with this, most ASVs are either unique to one quadrat or shared between the quadrats at the same site (Fig. 4**c**).

Few ASVs are present (at more than 3% of sample reads) in a high percentage of samples, showing a limited core cyanobacterial community with substantial variation between nearby hosts and soils, even on the scale of a few centimeters between collection points. Across the whole dataset, the most common ASV (ASV7) only appeared in 35.25% of samples and most ASVs appeared in less than 10% (Table S4). ASV7 is similarly common when looking across all soils (36.85%) and hornworts (34.52%). When soil samples from individual sites are considered, there are larger core cyanobacterial communities. At Grossman Pond, only four ASVs are present in over half of samples and six are in between a quarter and a half. At Potato Hill, one

ASV (ASV7) is in slightly over half the samples and five other ASVs, all distinct from those in the Grossman Pond core, are in the quarter to half of samples range. Plants at a site had less consistent core sets of cyanobacteria than soils. At Grossman Pond, only four ASVs (10, 14, 9, and 120; bolded ASVs in Fig. 3) were in 25-50% of plant samples and at Potato Hill, there were five in this category (7, 4, 2, 3, and 5). In summary, cyanobacteria appear to exhibit high heterogeneities in both soil and plant samples.

### Time is not a significant organizing factor in hornwort and soil cyanobacteria communities

Overall we found that sampling time explained the least of the community variance (R^2^=0.031, *p*<0.001) compared to location and sample type (Table S3). When each quadrat was analyzed separately, we also did not see samples clustered by time points on the PCoA plots (Fig. S5) and PERMANOVA tests found no significant association. The only exception is quadrat 1 in Potato Hill (*p*<10^−4^), which is largely driven by the *Notothylas* samples that did show a time-structured pattern (Fig. S5**c**). Our ANCOM analysis could not identify any individual ASV that consistently changes across the sampling time points within each quadrat. ASVs that appear in all time points or only one time point are the most common categories across all quadrats at both sites (Fig. S6).

### Cyanobiont communities are not strongly host specific but hosts show some selectivity

The three sympatric hornwort species co-occurring at Potato Hill (Fig. 1**c**) offer an ideal case to examine specificity (partner compatibility) and selectivity (partner preference) between hosts and cyanobionts. We found no difference in alpha diversity of overall cyanobiont communities of the three hosts (Table S5). Together with the large numbers of shared cyanobionts (Fig. 3), this indicates that the hosts were similar in specificity.

In terms of selectivity, if there are differences in partner preferences (i.e. certain hornwort species prefer certain cyanobacteria, or vice versa), we would expect to see cyanobiont communities cluster by host identity. We do not observe this in ordinations of all samples or only Potato Hill plants, but see some signal when looking at individual quadrats. A by-host association was found in quadrat 3 (PERMANOVA *p* < 10^−4^, R^2^=0.352; Fig. 5**c**) with *Notothylas* and *Anthoceros* samples mostly separating along the first PCoA axis (Fig. 5**e**). A difference also appeared in quadrat 2 based on PERMANOVA (*p* = 0.023, R^2^=0.126; Fig. 5**c**,**f**), but no difference in partner preference can be found in quadrat 1 (*p*=0.230).

**Figure 5.**
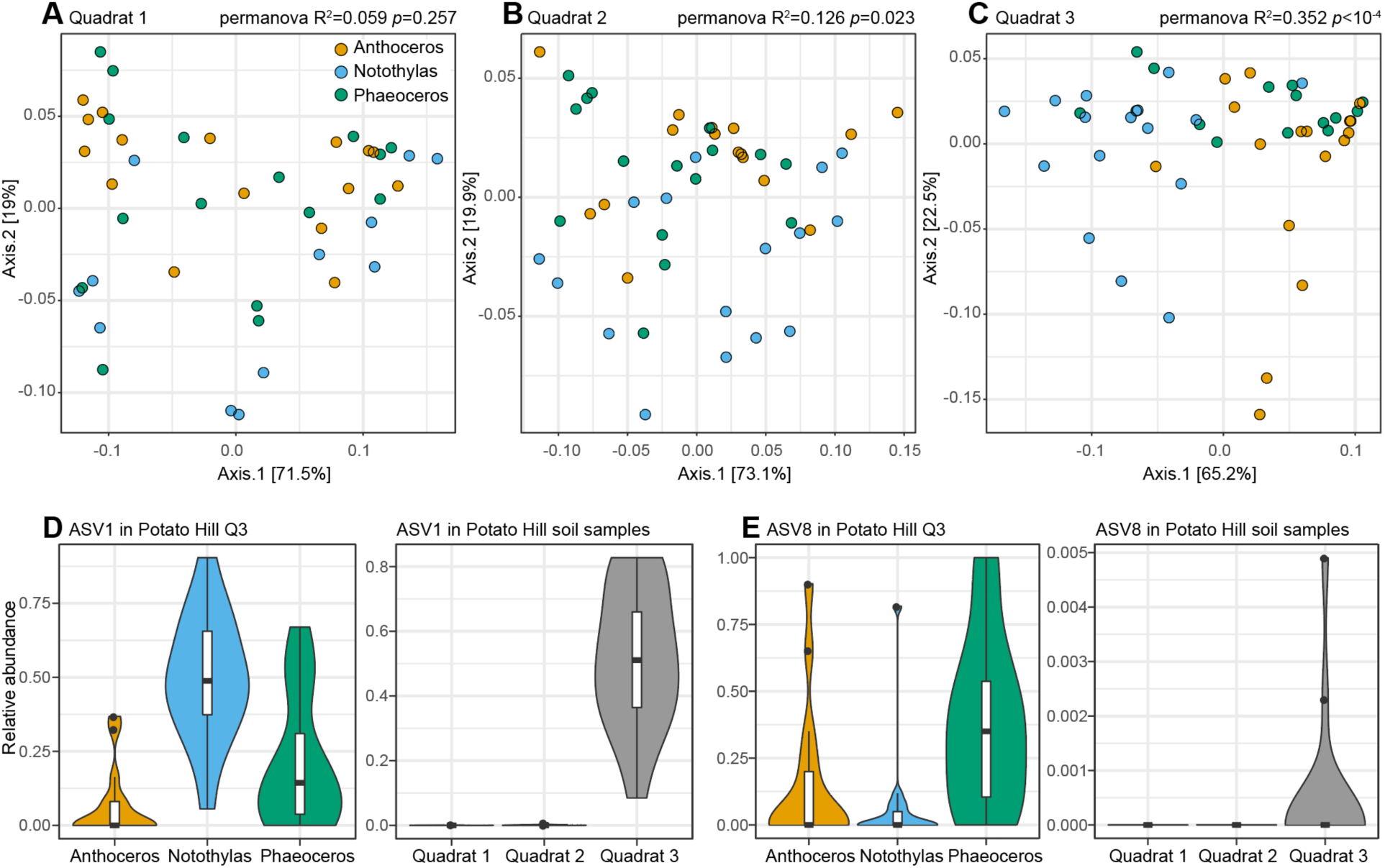
Context-dependent host-cyanobiont selectivity. PCoA plots (based on weighted unifrac distance) of (A) Quadrat 1, (B) Quadrat 2, and (C) Quadrat 3 in the Potato Hill site. The cyanobiont communities are color-coded by the host species, and permanova tests were done with host species as the grouping term. In Quadrat 3, cyanobiont communities are significantly organized by host species, with two ASVs that significantly vary in abundance among the three host species: (D) ASV1 and (E) ASV8 (the corresponding abundances in soil are shown on the right in each panel).

For quadrats 2 and 3, we further searched for the cyanobacteria ASVs that are differentially recruited by the three host species. Our ANCOM analyses identified ASV1 and ASV8 in quadrat 3, but none in quadrat 2, which is consistent with quadrat 2’s much weaker by-host interaction. ASV1 and ASV8 belong to two distantly related cyanobacteria clades (group I and *Nostoc* Subclade 3, respectively; Fig. 3) and exhibit different host association patterns:

ASV1 tends to have higher abundances in *Notothylas* samples (Fig. 5**d**), whereas ASV8 is more frequent in *Anthoceros* and *Phaeoceros* (Fig. 5**e**). These two ASVs are either absent or in very low abundance in quadrats 1 and 2 soil samples (Fig. 5**d**,**e**), which could explain, at least in part, why we observed little or no selectivity in those quadrats.

## Discussion

### Phylogenetic diversity of cyanobionts

Our data reveal that hornworts can form symbioses with phylogenetically diverse cyanobacteria within the order Nostocales, i.e. subclade B1 from Shih et al. (2013). Many hornwort cyanobionts are closely related to those from cyanolichens (e.g. *Peltigera*) or other plant associations. The overlaps between lichen and plant cyanobionts suggests that diverse fungal and plant lineages have independently tapped into the same cyanobacteria lineages for symbiotic partners, consistent with earlier studies based on fewer samples (Costa *et al*., 2001; O’Brien *et al*., 2005). Two lichen photobiont lineages, “*Nostoc* Subclade 2” and *Rhizonema* were not recovered in this study. However, this could simply reflect our geographic focus in temperate North America. “*Nostoc* Subclade 2” is mostly found in polar and boreal regions (Magain *et al*., 2017) and *Rhizonema* in the tropics (Lücking *et al*., 2009).

On the other hand, there are two cases where hornwort cyanobionts are either the only members of a clade (group II), or made up the bulk of a clade with a handful of free-living strains (group I; Fig. 3). Group I is particularly interesting. This clade diverged very deeply within Subclade B1 (Fig. 3), and consists of cyanobionts from all three hornwort species from Potato Hill (absent in Grossman Pond), as well as free-living cyanobacteria strains of “*Scytonema hofmanni*” UTEX 2349 (from Watkins Glen, New York) and “*Hassallia sp*” (from Mojave National Preserve, California). These two free-living strains each share an identical, or near-identical, *rbcL-X* sequence with at least another cyanobiont ASV. While some lichen-forming basidiomycetes were thought to harbor *Scytonema*, Lücking et al. (2009) showed that their photobionts are phylogenetically distinct from *Scytonema* and belong to a separate monophyletic genus *Rhizonema*. Group I is neither related to *Scytonema sensu stricto* nor *Rhizonema* (Fig. S2). In addition, the species identity of “*S. hofmanni*” UTEX 2349 is questionable, as it did not group with the *Scytonema* species nor with other *S. hofmanni* isolates. The other known strain in this clade was attributed to *Hassallia*, which has only one *rbcL-X* sequence available on Genbank and more data are needed to resolve its taxonomic status.

Overall, the broad diversity of cyanobionts even within our two sampling sites points to a wide selection of possible partners being compatible with hornworts, some of which probably belong to previously undescribed cyanobacterial lineages.

### Soil connection

Of all the studies on cyanobiont diversity in plants, only a few have examined the free- living cyanobacteria communities in the same habitats (Leizerovich *et al*., 1990; West & Adams, 1997; Rikkinen & Virtanen, 2008; Cuddy *et al*., 2012; Liaimer *et al*., 2016) and only one has done so using metabarcoding with next generation sequencing (Suárez-Moo *et al*., 2019).

Without knowing the background cyanobacteria compositions, it is difficult to infer and compare selectivity and specificity across species, or to understand the factors structuring cyanobiont communities. Our study is the first to comprehensively profile soil cyanobacteria on the same substrate as hornworts. We found that soil and hornwort cyanobiont communities overlapped significantly, contrary to the rather minor overlap found in previous work with cycads (Cuddy *et al*., 2012; Suárez-Moo *et al*., 2019) and bryophytes (West & Adams, 1997; Rikkinen & Virtanen, 2008; Liaimer *et al*., 2016). Such difference could be due to the fact that most earlier studies were culture-based and hence could only catalog a small portion of the soil diversity. More recently, Suarez-Moo et al. (2019) used 16S amplicon-seq to profile the microbiomes of coralloid roots of six Mexican cycad species (*Dioon spp*), as well as the immediate rhizosphere and bulk soils. Out of the 12 cyanobiont OTUs recovered, only 5 were also found in rhizosphere and/or soil samples. A similar study was done by Zheng and Gong (2019) on a Chinese *Cycas* species; however, very few cyanobacteria reads were recovered from their root and soil samples, making it difficult to compare the results. Our study suggests that hornworts might have a tighter soil connection than other plant lineages, although more amplicon studies are needed to validate this pattern. This difference could arise from the small, flat growth form of hornworts that results in a much higher percent of their surface being in direct contact with the substrate than is the case for larger, non- thalloid plants.

### Variation of cyanobacterial communities across time and space

Our sampling did not detect clear patterns of cyanobacterial community change over the time scale we surveyed. Based on the fact that the majority of the ASVs were either found in all time points or only one time point, it seems that some parts of a site’s cyanobacterial community is quite stable over the growing season, but this background is punctuated by rarer strains that only appear briefly. In the temperate habitat with long winters that we surveyed, it is likely that the community resets each year and future studies could investigate interannual variation.

The only other study using metabarcoding on hornwort bacteria (focusing on the tropical *Leiosporoceros dussii* which is unique in hosting cyanobacteria in channels rather than discrete colonies) found distinct communities between sites a few kilometers apart and the authors hypothesize that this could be due to differences in soil type (Bouchard *et al*., 2020). Our data confirm that the cyanobacteria can vary significantly not only between sites separated by a few kilometers but also even a few meters. Unsurprisingly, the distinction between soil communities is strongest between the two sites and less within sites. Plant samples from the two sites show less community separation, suggesting that even plants far from each other can obtain phylogenetically similar cyanobiont communities. Even so, even relatively small distances clearly have in impact since the Potato Hill plants cluster based on quadrat when only presence and absence of ASVs is considered (unweighted unifrac in Fig. S4).

In addition, we found that hornwort plants growing only a few centimeters apart can have very different sets of strains. Our study is the first to apply next-generation sequencing on hornworts that form discrete cyanobacterial colonies, confirming the within- and between-thallus variability of cyanobionts detected with culturing previously (West & Adams, 1997; Costa *et al*., 2001). Our data are consistent with separate slime cavities being independently colonized as they form (Adams & Duggan, 2008), but also show that individual hornworts are often dominated by one or two cyanobionts (Fig. S2). However, these dominant taxa differ between nearby plants, suggesting stochasticity in the colonization process. Future laboratory culture experiments could determine how much this variability may be due to competition between potential partners, priority effects, or random chance. Based on the high variability at small spatial scales, it will also be important to test in future work if the 0.25 m^2^ quadrat size can meaningfully represent hornwort cyanobiont communities.

### Specificity and selectivity

Sympatric species offer a unique opportunity to infer specificity and selectivity because the hosts are exposed to the same pool of symbionts (and vice versa). In our Potato Hill site, where *Anthoceros, Phaeoceros*, and *Notothylas* grow in sympatry, a large number of phylogenetically diverse ASVs are shared among the three host species (Fig. 3, Fig. 4**d**) and alpha diversity indices are similar across the hosts (Table S5). These results imply low reciprocal specificity between hornworts and cyanobacteria. Although Fig. 3 showed a few distinct ASVs that only came from *Notothylas*, they are restricted to the Grossman Pond site (dashed blue circles in Fig. 3) and absent in the soil samples elsewhere. Our data therefore indicate that the hornwort-cyanobacteria symbiosis is rather generalist, in contrast to the patterns of specificity observed for rhizobial and some mycorrhizal symbioses (Molina *et al*., 1992; Wang *et al*., 2012).

Some other plant lineages appear to have a higher specificity toward cyanobacteria than what we detect here for temperate hornworts. The most obvious case is *Azolla* for which the cyanobionts are known to be obligate and vertically transmitted (Zheng *et al*., 2009). A recent phylogenomic study demonstrated a strict one-to-one relationship between the two partners (Li *et al*., 2018). On the other hand, cycads do not have obligate cyanobionts, but the latest amplicon-seq results suggest that the interaction might be more specialized than what we found in hornworts. For example, cyanobionts from Mexican *Dioon* species fell into a single clade within Nostocales (Suárez-Moo *et al*., 2019). No comparable amplicon-seq studies exist for *Gunnera* or *Blasia* cyanobionts and clearer cyanobacterial taxonomy would be necessary to clearly define the breadth of cyanobiont diversity that can associate with various hosts.

While the ranges of possible cyanobionts may differ between hosts, previous work has found little evidence of correlation between host identity and cyanobiont community (Gehringer *et al*., 2010; Gutiérrez-García *et al*., 2019; Suárez-Moo *et al*., 2019). We did not observe host species to be a strong organizing factor, but our sympatric sampling hinted at some differential selectivity of the three hornwort host species. In particular, in quadrat 3 in Potato Hill, cyanobacteria communities are significantly clustered by host species (Fig. 5**c**). ANCOM analysis further identified two ASVs that are differentially abundant among the three hornwort species (Fig. 5**d**,**e**). Because these two ASVs are absent in soils of other quadrats (Fig. 5**d**,**e**), this selectivity is apparently dependent on the background cyanobacteria pool. Further laboratory experiments are needed to determine the direction of selection—whether it was hornworts selecting cyanobacteria or cyanobacteria selecting hornworts—and to explore if competition for partners (and not selection per se) can also result in the by-host association observed here.

## Conclusion

This study opens new opportunities for understanding the ecology and natural history of plant-cyanobacteria symbiosis. We demonstrated the efficacy of *rbcL-X* PacBio metabarcoding approach to profile cyanobacteria communities. With this method, we uncovered a high diversity of hornwort cyanobionts, some of which were not closely related to lichen photobionts nor other known plant-associated cyanobacteria. Three sympatric hornwort species have similarly low specificity toward cyanobionts, but can exhibit a significant difference in preferences among potential partner strains depending what is available in the soils. The soil connection appears to be tighter in hornworts than other plant-cyanobacteria symbiosis, although comparable studies are limited. Finally, we showed that the cyanobiont communities were not structured by sampling time points, but by distance at various scales. Our results highlight the importance of sampling soil and sympatric species to tackle the drivers of symbiotic community composition and mark an important step for using hornworts to explore these dynamics.

## Acknowledgments

This work was supported by a National Science Foundation grant (DEB-1831428) to F.-W. Li. We thank Norm Trigoboff for the information on hornwort distribution in Potato Hill State Forest, and Juan Carlos Villarreal for invaluable comments.

## Author contributions

J.M.N. and F.-W.L. conceived the project, carried out the sampling and data analyses, and wrote the manuscript. J.M.N. and D.A.H. carried out the molecular work.

## Supplementary Information

Figure S1. **Assessment of sufficient sequencing coverage**.

Figure S2. **Abundance distribution of the top ASV per sample**.

Figure S3. **Phylogeny of ASVs from this study and other *rbcL-X* sequences**.

Figure S4. **Cyanobacteria community in the Potato Hill site**.

Figure S5. **Cyanobacteria community through time**.

Figure S6. **Shared and unique ASVs across the time points (T1-T4)**.

Table S1. **The barcode sequences used in PCR primers**.

Table S2. **PCR recipe**.

Table S3. **PERMANOVA table**.

Table S4. **Statistics on the core cyanobacteria in different datasets**.

Table S5. **Comparison of cyanobacteria alpha diversity indices among the sampled species**.

