## Supplementary material for "Symbiotic cyanobacteria communities in hornworts across time, space, and host species": Figure S1

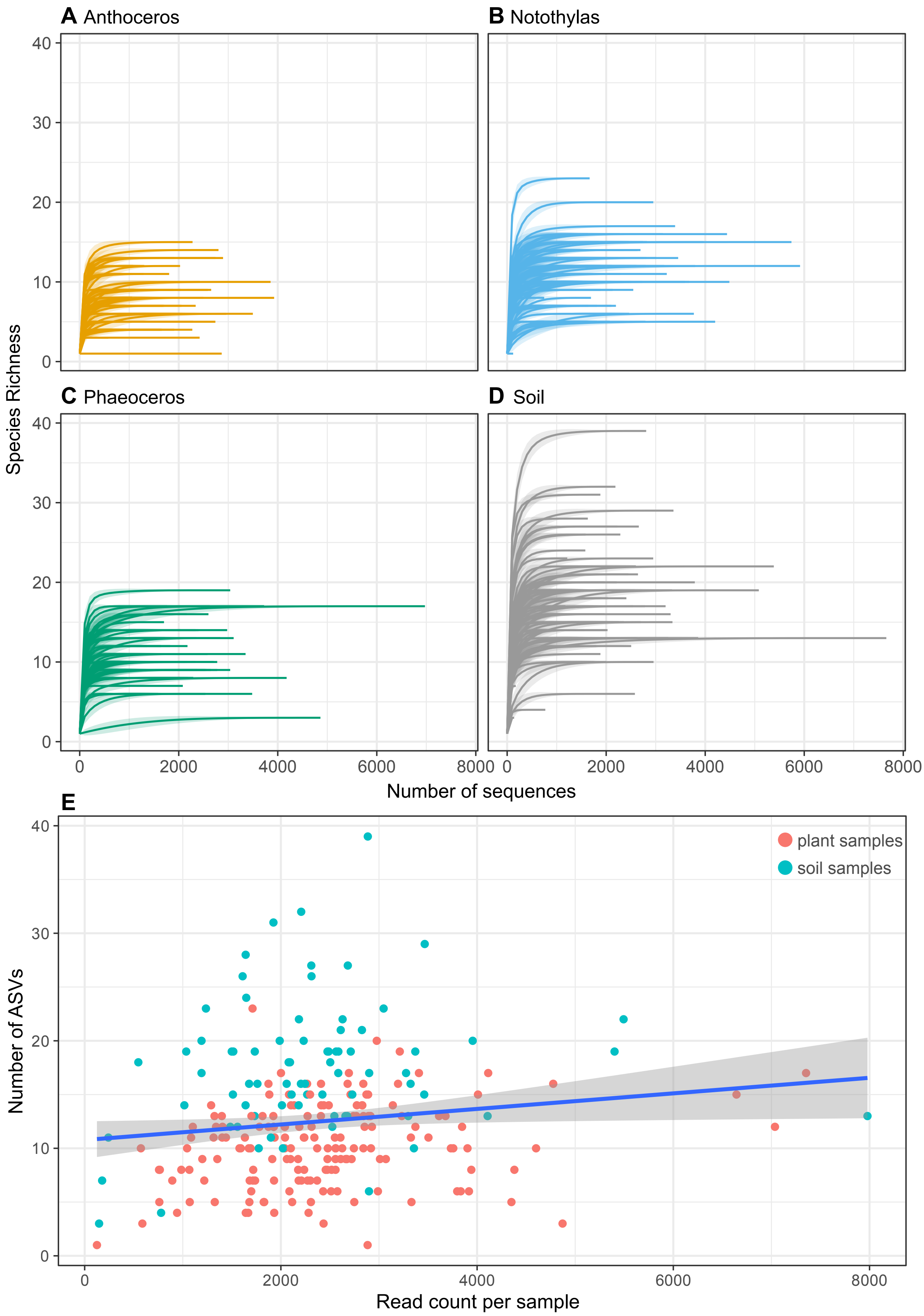


Supplementary Figure 1. **Assessment of sufficient sequencing coverage.** Rarefaction curves of (A) Anthoceros samples, (B) Notothylas samples, (C) Phaeoceros samples, and (D) soil samples all plateaued. (E) No strong correlation was found between ASV and read counts across all samples, which is consist with sufficient sequencing coverage. Linear regression line is shown in blue, with confidence interval in grey.
