## Supplementary material for "Symbiotic cyanobacteria communities in hornworts across time, space, and host species": Figure S2

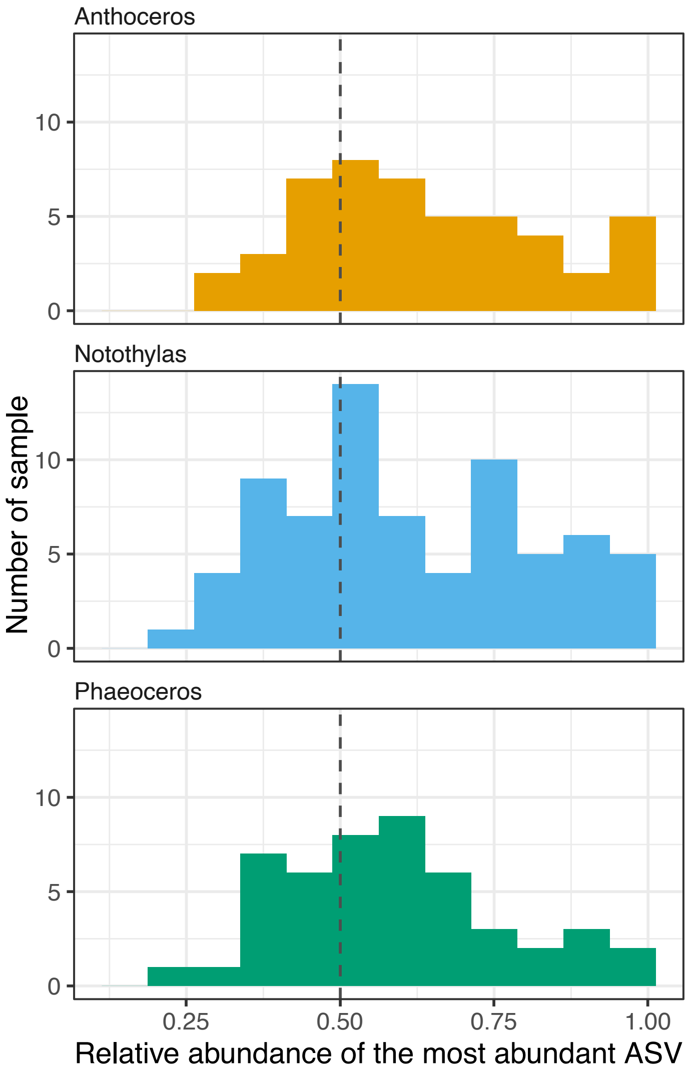


Supplementary Figure 2. **Abundance distribution of the top ASV per sample.** The dash line marks the 50% relative abundance.
