## Supplementary figures and images for "Symbiotic cyanobacteria communities in hornworts across time, space, and host species"

### Figure S3

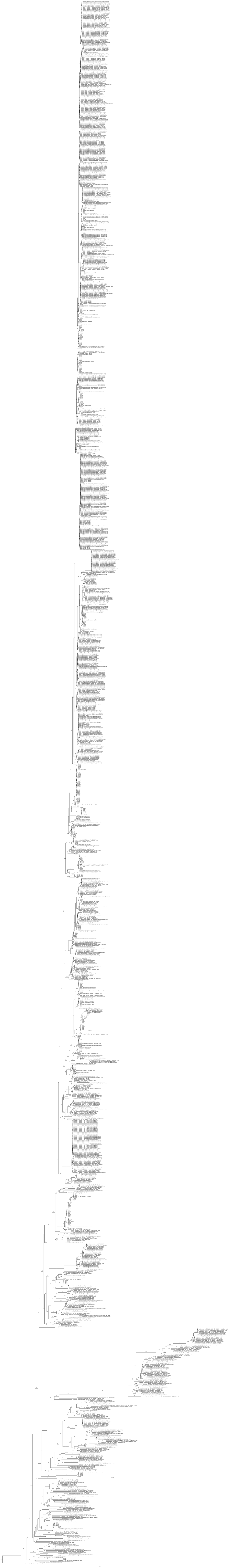

### Figure S6

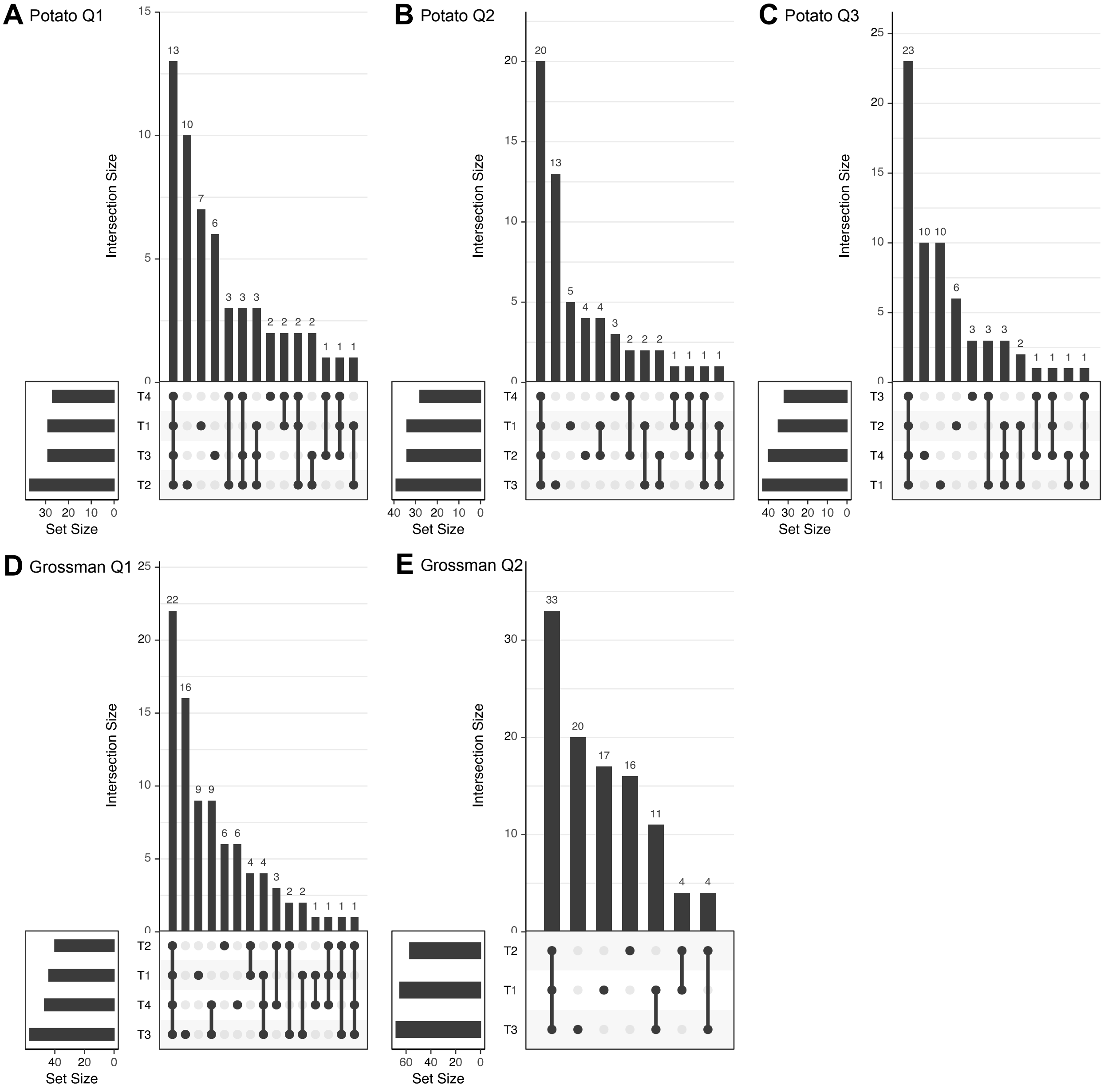


Supplementary Figure 6. **Shared and unique ASVs across the time points (T1-T4).**
