## Supplementary material for "Symbiotic cyanobacteria communities in hornworts across time, space, and host species": Figure S4

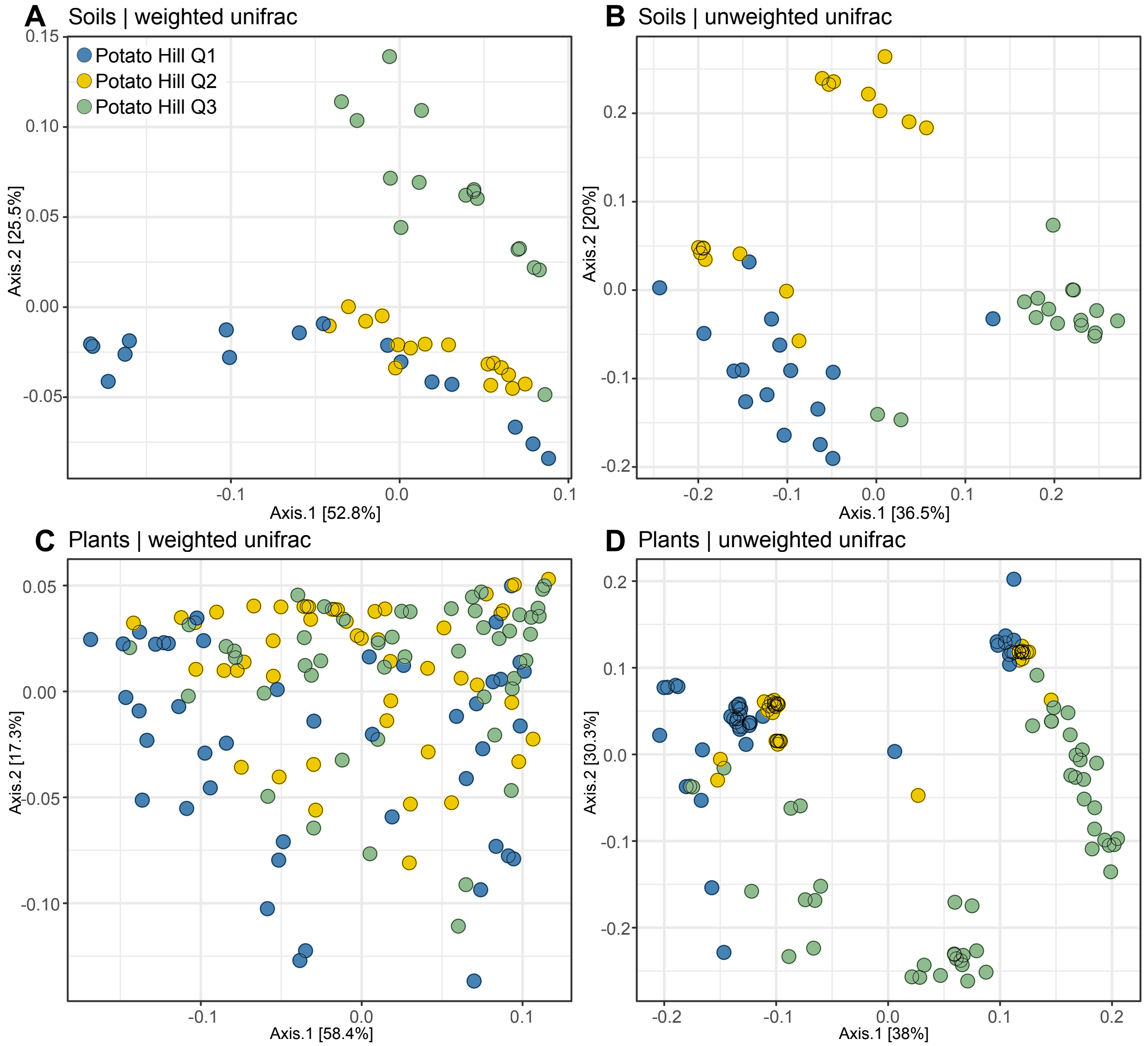


Supplementary Figure 4. **Cyanobacteria community in the Potato Hill site.** PCoA plots of the soil samples based on (A) weighted unifrac distance and (B) unweighted unifract distance, and plant samples based on (C) weighted unifrac distance and (D) unweighted unifract distance.
