## Supplementary material for "Symbiotic cyanobacteria communities in hornworts across time, space, and host species": Figure S5

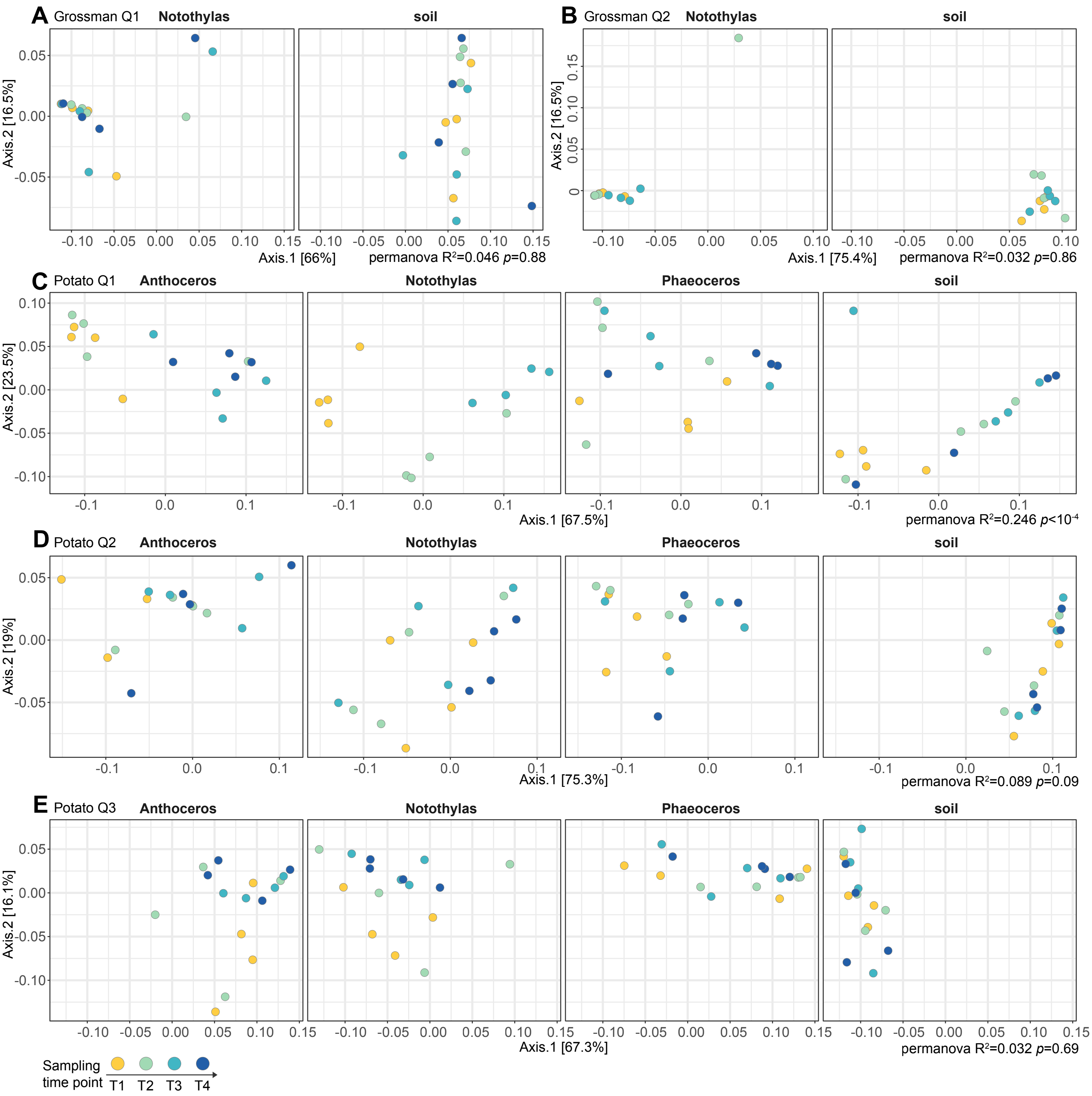


Supplementary Figure 5. **Cyanobacteria community through time.** PCoA plots based on weighted unifrac distance of (A) Grossman Pond Quadrat 1, (B) Grossman Pond Quadrat 2, (C) Potato Hill Quadrat 1, (D) Potato Hill Quadrat 2, (E) Potato Hill Quadrat 3. Samples were color-coded by sampling time points, and different sample types split into separate panels (but on the same ordinations). Permanova results were shown on the lower right for each quadrat (with time point as the grouping term).
